# *Fasciola hepatica* juveniles interact with the host fibrinolytic system as a potential early-stage invasion mechanism

**DOI:** 10.1101/2022.11.07.515559

**Authors:** Judit Serrat, David Becerro-Recio, María Torres-Valle, Fernando Simón, María Adela Valero, María Dolores Bargues, Santiago Mas-Coma, Mar Siles-Lucas, Javier González-Miguel

**Affiliations:** Laboratory of Parasitology, Institute of Natural Resources and Agrobiology of Salamanca (IRNASA-CSIC), 37008 Salamanca, Spain; Laboratory of Parasitology, Faculty of Pharmacy, University of Salamanca, 37007 Salamanca, Spain; Departamento de Parasitología, Facultad de Farmacia, Universidad de Valencia, Av. Vicent Andres Estelles s/n, 46100 Burjassot, Valencia, Spain; CIBER de Enfermedades Infecciosas, Instituto de Salud Carlos IIII, C/ Monforte de Lemos 3-5. Pabellón 11. Planta 0, 28029 Madrid, Spain

## Abstract

**Background:** The trematode *Fasciola hepatica* is the most widespread causative agent of fasciolosis, a parasitic disease that mainly affects humans and ruminants worldwide. During *F. hepatica* infection, newly excysted juveniles (FhNEJ) emerge in the duodenum of the mammalian host and migrate towards the definitive location of the parasite, the intra-hepatic biliary ducts. Understanding how *F. hepatica* traverses the intestinal wall and migrates towards the liver is pivotal for the development of more successful strategies against fasciolosis. The central enzyme of the mammalian fibrinolytic system is plasmin, a serine protease whose functions are exploited by a number of parasite species owing to its broad spectrum of substrates, including components of tissue extracellular matrices. The aim of the present work is to understand whether FhNEJ co-opt the functions of their host fibrinolytic system as a mechanism to facilitate trans-intestinal migration.

**Methodology/Principal Findings:** An FhNEJ tegument protein extract (FhNEJ-Teg) was obtained *in vitro*, and its capability to bind the zymogen plasminogen (PLG) and enhance its conversion to the active protease, plasmin, were analyzed by a combination of enzyme-linked immunosorbent, chromogenic and immunofluorescence assays. Additionally, PLG-binding proteins in FhNEJ-Teg were identified by 2D electrophoresis coupled to mass-spectrometry analysis, and the interactions were validated using FhNEJ recombinant proteins.

**Conclusions/Significance:** Our results show that FhNEJ-Teg contains proteins that bind PLG and stimulate its activation to plasmin, which could facilitate the traversal of the intestinal wall by FhNEJ and contribute to the successful establishment of the parasite within its mammalian host. Altogether, our findings contribute to a better understanding of host-parasite relationships during early fasciolosis and may be exploited from a pharmacological and/or immunological perspective for the development of treatment and control strategies against this global disease.

**Author Summary:** Fasciolosis is a disease caused by parasites of the genus *Fasciola*, of which *F. hepatica* stands out as it has successfully spread all over the world and infects humans and animals throughout the entire global geography. Definitive hosts become infected by ingestion of aquatic plants or water contaminated with metacercariae, which excyst in the duodenum and release the so-called newly excysted juvenile flukes (FhNEJ). FhNEJ traverse the intestinal wall and evolve into immature parasites that migrate through the peritoneum and liver parenchyma until they reach their definitive location inside the major biliary ducts, where adult worms develop and egg shedding starts. In order to cross the intestinal wall, FhNEJ are endowed with a repertoire of proteases that degrade components of the intestinal extracellular matrix, and we hypothesized that they may also co-opt the proteolytic functions of plasmin, the central enzyme of the mammalian fibrinolytic system, to migrate more efficiently across host tissues. In this study, we demonstrate that FhNEJ express proteins on their tegument surface that interact with plasminogen, the zymogen of plasmin, and stimulate its conversion into its active form, which could potentially be used for trans-intestinal migration and contribute to the successful establishment of the parasite within its mammalian host.

## Introduction

Fasciolosis caused by the trematode *Fasciola hepatica* is considered the widest-spread helminth disease, affecting wild and domestic animals and up to 17 million people worldwide (1). Definitive hosts (typically ruminants raised as livestock and humans) become infected by ingestion of metacercariae commonly attached to water plants that are used for consumption. Upon ingestion, metacercariae excyst in the duodenum and release the newly excysted juvenile flukes (FhNEJ), which cross the intestinal wall within the first two to three hours of infection and migrate throughout the peritoneum towards the liver. After burrowing through and feeding from hepatic tissue, migrating juveniles finally establish in the major biliary ducts, where they mature into adult worms and egg shedding starts (2). Provided that human fasciolosis hyper-endemic areas are found in regions where people live in substandard socio-economic conditions (2–4), this parasitosis is also classified by the World Health Organization as a Neglected Tropical Disease (5). Nowadays, the emergence of resistant isolates against the drug of choice triclabendazole, the lack of an effective vaccine against *F. hepatica* and the geographic expansion of this parasite due to climate change turns fasciolosis into a disease of growing public health concern (6,7).

Classical vaccine strategies against *F. hepatica* have focused on targeting the adult phase of this parasite, which leads to some drawbacks considering that i) migrating juveniles produce substantial tissue damage during the acute stage of the disease (8), which occasionally causes sudden death of the definitive host (9), and ii) adult flukes reside in the liver, an anatomical niche that preferentially induces immunotolerance. In this scenario, interventions aimed at blocking the migration of juveniles and the development of adult flukes may be more effective in fasciolosis prevention and control, but the molecular mechanisms involved in FhNEJ survival and development inside the host are not yet fully understood. Given that crossing the intestinal wall by FhNEJ can be regarded as a ‘point of no return’ in fasciolosis in terms of therapeutic control (8), it is of utmost importance to precisely understand the molecular events that regulate this process.

Crossing of the intestinal wall by FhNEJ is driven, among others, by cathepsin proteases, which are papain-like cysteine peptidases that are capable of degrading components of the intestinal extracellular matrix (ECM) (10,11). *F. hepatica* cathepsins are amongst the most highly expressed proteins in *F. hepatica*, which evidences the importance of these enzymes on *F. hepatica* biology and development, and they are divided in two families (B and L) that show a temporal pattern of expression, being FhCL3 and FhCB1, FhCB2 and FhCB3 the most highly expressed isotypes in FhNEJ (12).

In addition to their endogenous repertoire of proteases, it is possible that FhNEJ co-opt proteolytic functions of the host in order to migrate more efficiently in terms of energy expenditure. A paradigmatic example of exploitation of host resources is the interaction between parasites and plasminogen (PLG), the central pro-enzyme of the mammalian fibrinolytic system (13–15), which has been described in adult *F. hepatica* via the secretion of PLG-binding proteins and proposed to be a mechanism of survival inside the biliary ducts (16,17). PLG is converted into its active form, plasmin, by the serine proteases tissue-type or urokinase-type plasminogen activators (t-PA and u-PA, respectively). PLG binding to fibrin and other partner proteins occurs via specialized protein domains found in the PLG molecule called kringle, which have high affinity for lysine residues of partner proteins (18). Although t-PA expression is mostly restricted to the vascular endothelium, u-PA is expressed in different tissues across the organism and can therefore stimulate plasmin generation in the extravascular space (19). This, in conjunction with the wide array of plasmin substrates, including ECM components (19), matrix metalloproteinases (20) and the C3 and C5 complement molecules (21), explains why host-derived plasmin could be a highly valuable tool for parasites to migrate through the mammalian organism, modulate immune responses and overall increase their chances of survival within their definitive hosts (13–15).

The aim of the present work is to investigate whether FhNEJ interact with the fibrinolytic system and promote plasmin generation at their surface as a potential mechanism to facilitate trans-intestinal migration. This study reveals the pro-fibrinolytic potential of FhNEJ and broadens our knowledge on host-parasite relationships during early infection by *F. hepatica*.

## Methods

### *In vitro* excystment of *F. hepatica* metacercariae

FhNEJ were obtained by *in vitro* stimulating the excystment of five thousand *F. hepatica* metacercariae (Ridgeway Research Ltd) as previously described (22). Briefly, pure CO_2_ was bubbled in 10 ml of ice-cold distilled water for 30 sec, followed by supplementation with sodium dithionite to a final concentration of 0.02 M. This solution was added to metacercariae and incubated for 1 hour at 37 °C. Following incubation, the metacercariae were washed twice with room-temperature distilled water and resuspended in 5 ml of Hank’s balanced salt solution (Sigma) supplemented with 10% lamb bile (obtained from a local abattoir) and 30 mM HEPES (Sigma) pH 7.4. The parasites were incubated for four hours at 37 °C and FhNEJ were manually recovered under a magnifier using a 20 μl pipette and immediately subjected to protein extraction.

### Protein extraction of the tegument fraction

After two washes in sterile phosphate-buffered saline (PBS), FhNEJ were resuspended in 300 μl of PBS containing 1% Nonidet P40 (NP40) and incubated at room temperature for 30 min with mild rotation. Next, parasites were centrifuged five min at 300 × *g* and the supernatant containing the FhNEJ tegument protein extract (FhNEJ-Teg) was frozen at −80 °C until use (23). Protein concentration was determined using the Pierce BCA Protein Assay kit (Thermo Fisher).

### Plasminogen binding assay

PLG binding was assessed by enzyme-linked immunosorbent assay (ELISA) as previously described (24). In brief, 96 well plates (Costar) were coated overnight at 4 °C with 0.5 μg of FhNEJ-Teg or 1 μg of recombinant protein diluted in carbonate buffer (15 mM Na_2_CO_3_, 35 mM NaHCO_3_, pH 9.6). Wells coated in 1% bovine serum albumin (BSA) were used as negative controls. After washing 3 times with PBS containing 0.05% Tween_20_ (PBST), wells were blocked with 1% BSA diluted in PBS for 30 min at 37 °C and increasing amounts (0 μg to 3 μg) of PLG (Origene) diluted in blocking solution were added to the wells and incubated for 1 hour at 37 °C. In parallel, a competition assay was performed in which 50 mM of 6-aminocaproic acid (ε-ACA) (Sigma) was added to the wells together with PLG. After 3 washes with PBST, the wells were incubated with an anti-PLG primary antibody raised in sheep (Acris Antibodies) followed by horseradish (HRP)-conjugated anti-sheep secondary antibody (Sigma). Primary and secondary antibodies were diluted 1:2000 and 1:4000, respectively, in blocking solution and incubated for 1 hour at 37 °C. Wells were washed 3 times with PBST between and after antibody incubations. Finally, PLG binding was revealed by adding 1.5 mM of the chromogenic substrate ortho-phenylene-diamine (OPD) (Sigma) diluted in substrate buffer (25 mM citric acid, 45 mM Na_2_HPO_4_, 0.04% H_2_O_2_, pH 5). The wells were incubated at room temperature in the dark and the reaction was stopped by adding an equal volume of 3 N sulfuric acid. Optical densities were measured at 492 nm in a Multiskan GO spectrophotometer (Thermo Fisher). The assay with FhNEJ-Teg was performed in duplicate, whereas those with recombinant cathepsins were performed in triplicate.

### Detection of plasminogen binding on the surface of FhNEJ by immunofluorescence

*F. hepatica* metacercariae were excysted as described above and FhNEJ were recovered every 1 hour after addition of excystment medium and cultured in RPMI-1640 culture media (Thermo Fisher) at 37 °C in a 5% CO_2_ atmosphere. Fifty FhNEJ per condition were washed 3 times in PBS and incubated with blocking solution (0.1% BSA in PBS) supplemented with 100 μg/ml of human PLG (Origene) for three hours at 37 °C. In parallel, a competition assay was performed that included 50 mM ε-ACA during human PLG activation. Two extra sets of FhNEJ incubated in blocking buffer alone served as negative controls for PLG staining and to control for unspecific background staining derived from the secondary antibody. After PLG incubation, FhNEJ were fixed in 4% paraformaldehyde (Santa Cruz) for 1 hour at room temperature and probed with 13 μg/ml of anti-human PLG antibody raised in sheep (Innovative Research) followed by incubation with Alexa Fluor 568 donkey anti-sheep IgG (Thermo Fisher) diluted 1:500. FhNEJ were washed 3 times with blocking buffer between incubations and primary and secondary antibodies were diluted in blocking solution and incubated overnight at 4 °C. Finally, FhNEJ were washed 3 last times in blocking solution and whole-mounted in glass slides using a 9:1 glycerol solution containing 0.1 M propyl gallate. Specimens were viewed using a spinning disk Dragonfly High Speed Confocal Microscope System (Andor, Oxford Instruments) located at the Microscopy Facility of the Institute of Functional Biology and Genomics (IBFG) (Salamanca, Spain) under the 40x/0.95 dry objective lens. A minimum of 10 FhNEJ were acquired per condition. Stacks of 30 slices (1 μm/slice) spanning the entire FhNEJ volume were acquired and images were processed in FIJI software (25) by getting a sum slices Z projection of each FhNEJ. For visualization purposes, histogram widths were equally adjusted in all conditions, images were converted to RGB color and exported as .tiff files.

### Plasminogen activation assay

This assay was performed in 96 well plates (Costar) in a final volume of 100 μl by measuring the amidolytic activity of generated plasmin on a plasmin-specific chromogenic substrate as previously described (24). In every well, 2 μg of PLG (Origene) were diluted in PBS together with 3 μg of D-Val-Leu-Lys 4-nitroanilide dihydrochloride chromogenic substrate (S-2251) (Sigma) and 1 μg of FhNEJ-Teg or recombinant protein in the presence or absence of 15 ng t-PA (Sigma) or 10 ng u-PA (Sigma). FhNEJ-Teg or recombinant proteins were replaced by equal amounts of BSA and incubated in the presence or absence of PLG activators to control for both the capability of t-PA or u-PA to activate unbound PLG and the spontaneous conversion of PLG into plasmin, respectively. Microplates were incubated for three hours at 37 °C and substrate cleavage was assessed by measuring absorbance at 405 nm every 30 min in a Multiskan GO spectrophotometer (Thermo Fisher). All the reactions were performed in triplicate.

### Detection of plasminogen activation on the surface of live FhNEJ

This assay is an adaptation of the PLG activation assay described in the previous section including incubation with live FhNEJ. FhNEJ were obtained by stimulating the excystment of *F. hepatica* metacercariae as previously described (see section “*In vitro* excystment of *F. hepatica* metacercariae”) and were left to recover in RPMI-1640 culture media (Thermo Fisher) for three hours at 37 °C in a 5% CO_2_ atmosphere after excystment. In every well, 2 μg of PLG (Origene) were diluted in PBS together with 3 μg of S-2251 (Sigma) and 20 FhNEJs in the presence or absence of 15 ng t-PA (Sigma) plus 10 ng u-PA (Sigma). Some wells contained FhNEJ-Teg in place of live FhNEJ as a positive control for plasmin generation. Wells containing PBS only, FhNEJ in PBS or PLG plus t-PA and u-PA (without FhNEJ) were used as negative controls. Microplates were incubated for three hours at 37 °C and substrate cleavage was assessed by measuring absorbance at 405 nm every 30 min in a Multiskan GO spectrophotometer (Thermo Fisher). All the reactions were performed in quadruplicate.

### Bidimensional (2D) electrophoresis of FhNEJ-Teg

First, FhNEJ-Teg extract was purified using the ReadyPrep 2-D Cleanup Kit (BioRad) and the protein pellets resuspended in rehydration buffer [7 M urea, 2 M thiourea, 4% 3-[(3-cholamidopropyl) dimethylammonio]-1-propanesulfonate (CHAPS)]. The extract was then divided in aliquots of 125 μl, supplemented with 0.05 M dithiothreitol (DTT) (Sigma) and 0.2% ampholytes pH 3-10 (BioRad) and added to 7 cm ReadyStrip IPG Strips with a linear pH range of 3-10 (BioRad) for passive rehydration overnight at 20 °C in a Protean IEF Cell equipment (BioRad). Isoelectric focusing (IEF) was performed the next day by using a constant amperage of 50 μA per strip in the Protean IEF Cell equipment (BioRad). Next, the proteins in the strips were reduced with DTT (0.02 g/ml) and alkylated with iodoacetamide (0.0025 g/ml) for 10 min and 15 min, respectively, at room temperature (both DTT and iodoacetamide were diluted in equilibration buffer containing 6M urea, 2% SDS, 1.5M Tris-HCl pH 8.8, 30% glycerol and bromophenol blue), and the separation for the second dimension was done in 12% acrylamide-sodium dodecyl sulfate (SDS) gels. These gels were stained with silver using in-house prepared reagents following the standard protocol (excluding formaldehyde and glutaraldehyde from the formulations to ensure compatibility with subsequent analysis by mass spectrometry) or transferred to a nitrocellulose membrane for immunoblot detection of PLG binding. Silver-stained gels were imaged using a Chemidoc gel-imaging system (BioRad).

### Immunoblot detection of plasminogen binding

The proteins in the 2D gels were transferred to nitrocellulose membranes using a constant amperage of 400 mA for 90 min at 4 °C. Blots were blocked for 1 hour at room temperature with 2% BSA diluted in PBST and incubated overnight at 4 °C with 25 μg/ml of PLG (Origene) diluted in blocking solution. PLG binding was detected by adding an anti-PLG primary antibody raised in sheep (Acris Antibodies) followed by horseradish (HRP)-conjugated anti-sheep secondary antibody (Sigma). Primary and secondary antibodies were diluted 1:1000 and 1:2000, respectively, in blocking solution and incubated for 90 min at 37 °C. Membranes were washed 3 times with PBST between incubations and protein spots with bound PLG were revealed with 4-chloro-naphtol following standard procedures. Blots were imaged in a Chemidoc gel-imaging system (BioRad) and spot matching between silver-stained gels and the blots and the assignment of molecular weights (MW) and isoelectric points (pI) of each protein were performed using the PDQuest Software v.8.0.1 (BioRad).

### Spot analysis by liquid chromatography coupled to tandem mass spectrometry (LC-MS/MS)

Protein spots in the silver stained gels were manually excised using 1000 μl sterile pipette tips and sent for proteomic analysis at the proteomics facility of the Central Support Service for Experimental Research (SCSIE, University of Valencia).

In-gel protein digestion of every spot was performed using sequencing-grade trypsin (Promega) as described elsewhere (26). In brief, 100 ng of trypsin were used for each sample, and digestion was performed overnight at 37 °C. Trypsin digestion was stopped with 10% trifluoroacetic acid (TFA) and the supernatant was removed. Next, samples were subjected to double acetonitrile (ACN) extraction and the peptide mixtures were dried in a speed vacuum and resuspended in 9 μl of 2 % ACN, 0.1% TFA.

LC-MS/MS was carried out as follows: 5 μl of digested peptide mixtures were loaded onto a trap column (3μ C18-CL, 350 um × 0.5mm) (Eksigent) and desalted with 0.1% TFA at 5 μl/min during five min. The peptides were then loaded onto an analytical column (3μ C18-CL 120 Å, 0.075 × 150 mm) (Eksigent) equilibrated in 5% ACN plus 0.1% formic acid (FA). Elution was carried out with a linear gradient of 7-40% B in A for 20 min (A: 0.1% FA; B: CAN plus 0.1% FA) at a flow rate of 300 nL/min. Peptides were analyzed in a mass spectrometer nanoESI qQTOF (6600plus TripleTOF) (ABSCIEX). Sample was ionized in a Source Type: Optiflow<1 μL Nano applying 3.0 kV to the spray emitter at 200 °C. Analysis was carried out in a data-dependent mode. Survey MS1 scans were acquired from 350–1400 m/z for 250 ms. The quadrupole resolution was set to ‘LOW’ for MS2 experiments, which were acquired 100-1500 m/z for 25 ms in ‘high sensitivity’ mode. The following switch criteria were used: charge 2+ to 4+, minimum intensity and 250 counts per sec. Up to 100 ions were selected for fragmentation after each survey scan. Dynamic exclusion was set to 15 sec. Finally, the rolling collision energies equations were set for all ions as for 2+ ions according to the following equations: |CE|=(slope)x(m/z)+(intercept).

### Protein identification

ProteinPilot v5.0 search engine (ABSCIEX) was used for protein identification. ProteinPilot default parameters were used to generate a peak list directly from 6600 plus TripleTOF wiff files. The Paragon algorithm (27) of ProteinPilot v5.0 was used to search the Uniprot_trematoda database (200604, 362615). The following parameters were used: Trypsin specificity, IAM cys-alkylation, no taxonomy restriction and the search effort set to rapid analysis. Protein grouping was done using the Pro Group algorithm (ABSCIEX). Only proteins belonging to *F. hepatica* and having an Unused value equal to or greater than 2 were used for subsequent analyses, and protein isoforms were manually grouped to facilitate downstream data interpretation. Gene Ontology analysis in the Biological Function category of PLG-binding proteins identified via 2D-MS was performed using the software Blast2GO v5.2.

### Statistical analysis

Plots were created with Prism 9 software (GraphPad Software, La Jolla, CA) and statistical analyses were performed with the R Commander package (28). Comparison between 2 groups used un unpaired Student’s *t*-Test. Comparison between 3 or more groups used an Analysis of Variance (ANOVA) test followed by a Tukey post-hoc analysis for pair-wise comparisons. Unless otherwise stated, the differences are not significant.

## Results

### The tegument of FhNEJ contains plasminogen-binding proteins

The ability of FhNEJ tegument proteins to bind PLG was analyzed by ELISA. To this end, wells were coated with FhNEJ-Teg, incubated with PLG, washed, and PLG binding was detected using standard downstream ELISA procedures. This experiment showed that FhNEJ-Teg contains proteins that are capable of binding PLG in a concentration-dependent manner as compared to the negative control, 1% BSA (Figure 1). In parallel to this, and aiming at underpinning the molecular mechanism of PLG interaction with parasite tegument proteins, a competition assay was performed in which the lysine analog ε-ACA was added to the reaction. In this case, PLG binding was completely abrogated, suggesting that PLG binding to FhNEJ-Teg is specific and mediated by its kringle domains.

**Figure 1.**
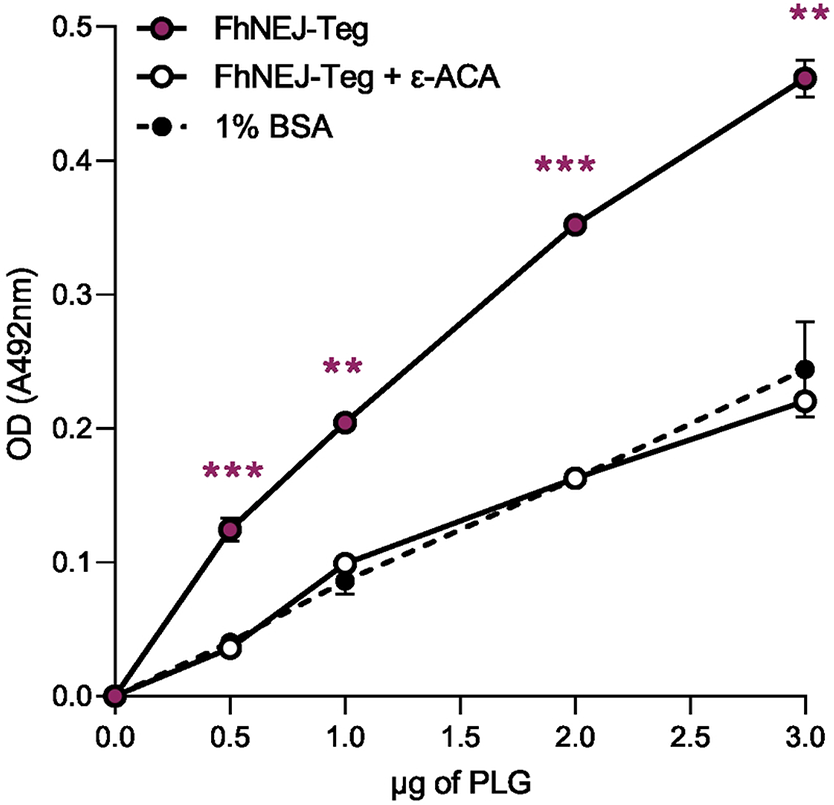
The tegument of FhNEJ contains proteins that bind plasminogen in a concentration- and lysine-dependent manner. FhNEJ-Teg binding to PLG was detected via ELISA by coating wells with 0.5 μg FhNEJ-Teg and incubating with increasing amounts of human PLG. In parallel, a competition assay was performed by including 50 mM of ε-ACA during PLG incubation. FhNEJ-Teg-coated wells and incubated with 1% BSA served as negative controls for PLG binding. Dots indicate the mean of two replicates ± SD. Asterisks indicate significant differences between FhNEJ-Teg and the rest of the groups (*p≤0.5; **p≤0.01; ***p≤0.001; one-way ANOVA followed by Tukey contrasts for pairwise comparisons).

In order to reveal the spatial distribution of PLG-binding proteins on the surface of FhNEJ, PLG binding was addressed via immunofluorescent staining by incubating FhNEJ in the presence or absence of PLG prior to fixation (Figure 2). Although we found some inter-individual variability in the staining pattern of FhNEJ incubated in the presence of PLG, we observed a trend of specific staining for PLG at the posterior area of FhNEJ, which most likely coincides with the excretory (protonephridial) pore (Figure 2B panel i). Additionally, some specimens also showed specific PLG staining at the oral sucker (Figure 2B panel ii) and some others all over the tegument surface (Figure 2B panel iii). FhNEJ incubated in the absence of PLG, in the presence of PLG together with 50 mM of ε-ACA or with secondary antibody alone (Figures 2A, 2C and 2D, respectively) had some unspecific background staining that was less intense than that observed in FhNEJ incubated in the presence of PLG.

**Figure 2.**
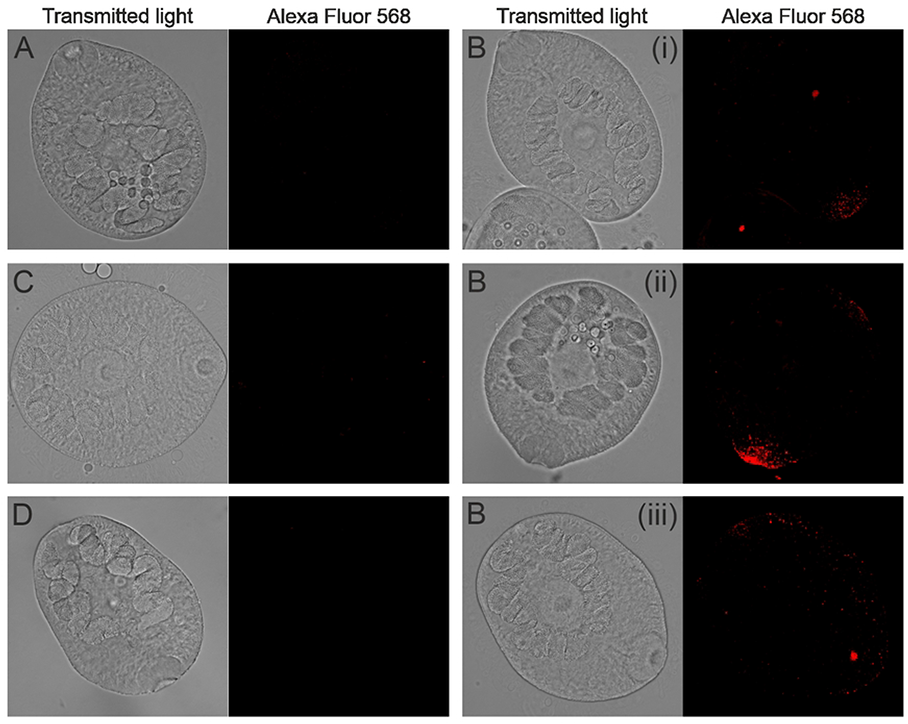
Detection of PLG binding at the surface of FhNEJ by immunofluorescence reveals distinct localization patterns of PLG-binding proteins. FhNEJ were whole-mounted and PLG binding (Alexa Fluor 568) was detected by immunolocalization using a confocal laser microscope. FhNEJs were incubated in the absence (A) or presence (B) of 30 μg/ml of PLG; in the presence of PLG plus 50 mM ε-ACA (C) or with secondary-antibody alone (D). PLG binding was only observed in FhNEJ incubated with PLG (B) and detected around the excretory (protonephridial) pore (B, panel i), the oral sucker (B, panel ii) and all over the tegument surface (B, panel iii).

### Binding of plasminogen by FhNEJ-Teg proteins enhances plasmin generation

In order to determine whether PLG binding by FhNEJ-Teg proteins facilitates the conversion of bound PLG to the catalytically active protease, plasmin, we performed an enzymatic assay by co-incubating FhNEJ-Teg with PLG and a plasmin-specific chromogenic substrate. This assay was performed in the presence or absence of the physiologic PLG activators, t-PA or u-PA, to investigate whether plasmin conversion from bound PLG is dependent on host-derived factors or FhNEJ-Teg contains proteases that are capable of cleaving and activating PLG themselves. As observed in Figure 3, the conversion of PLG to plasmin by both t-PA (Figure 3A) and u-PA (Figure 3B) is enhanced in the presence of FhNEJ tegument proteins.

**Figure 3.**
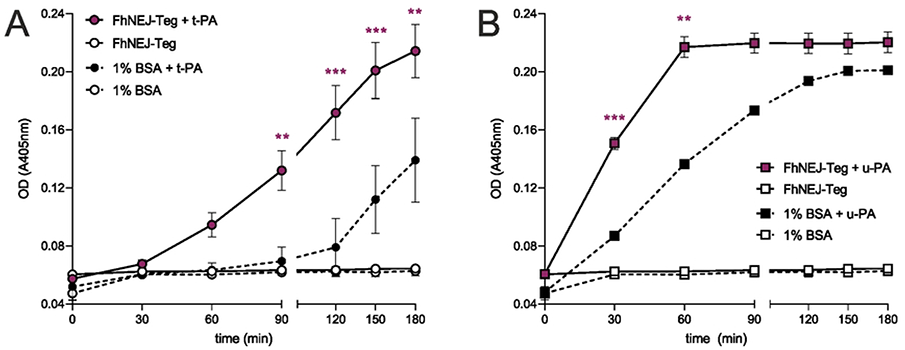
Plasminogen binding by FhNEJ tegument proteins facilitates its conversion to plasmin. One μg of FhNEJ-TEG was incubated with human PLG, a chromogenic substrate for plasmin (S-2251) and either t-PA (A) or u-PA (B) and plasmin generation was assessed by measuring substrate cleavage at OD 405 nm. In some instances, FhNEJ-Teg was replaced by 1% BSA as a negative control. Dots indicate the mean of three replicates ± SD, and asterisks indicate significant differences between FhNEJ-Teg + t-PA (A) or u-PA (B) and its BSA + PA counterpart (*p≤0.05, **p≤0.001, ***p≤0.001, n.s. not significant; one-way ANOVA followed by Tukey contrasts for pairwise comparisons).

We next investigated whether plasmin generation also occurs in the surface of live FhNEJ, so we set up a similar enzymatic assay to test for plasmin activation using live parasites next to the FhNEJ-Teg extract (Figure 4). We confirmed that FhNEJ were alive throughout the entire experiment by observing under a stereoscope that they remained highly mobile. Twenty FhNEJ per well were used since based on our expertise this amount of FhNEJs should contain approximately 1 μg of tegument proteins, which is the amount that we employed in the abovementioned enzymatic assay. Remarkably, we observed that live FhNEJ incubated with PLG are capable of enhancing plasmin generation in the presence of PLG activators, and that this effect is significantly higher than that observed in wells containing PLG and its activators without FhNEJ. Additionally, the potentiation of plasmin generation by 20 live FhNEJ almost identically replicates the effects obtained when 1 μg of FhNEJ-Teg is added to the reaction (Figure 4).

**Figure 4.**
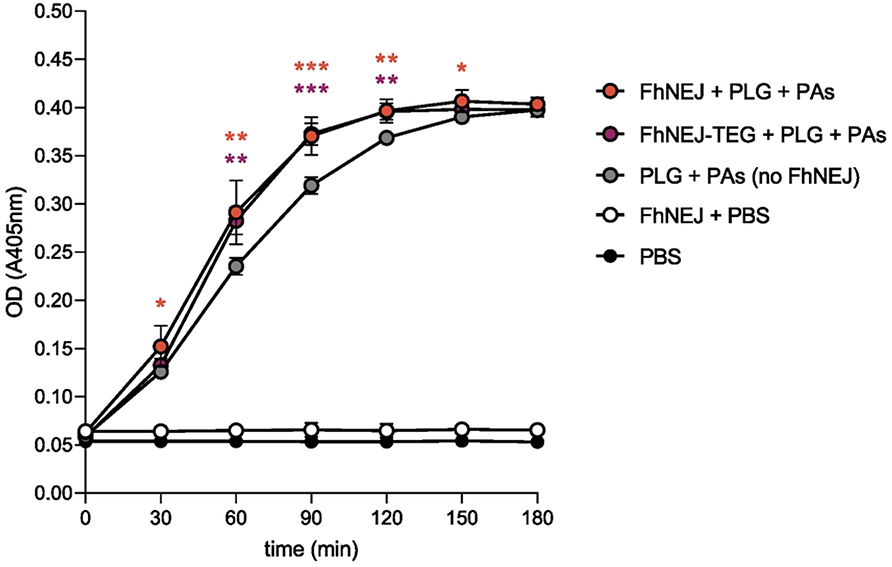
Detection of plasmin generation on the surface of live FhNEJ. Twenty FhNEJ per well were incubated with PLG, PLG activators (PAs) and a chromogenic substrate for Plm (S-2251), and the capability of live FhNEJ to stimulate plasmin generation on their surface was determined by measuring substrate cleavage (absorbance at 405 nm) every 30 minutes. Dots indicate the mean of four biological replicates ± SD. Wells containing PBS, FhNEJ in PBS and PLG + PAs without FhNEJ serve as negative controls; and wells containing PLG + PAs + FhNEJ-Teg (1 μg per well) serve as a positive control for the stimulation of plasmin generation by FhNEJ-derived proteins. Upper asterisks indicate significant differences between the negative control PLG + PAs without FhNEJ and its counterpart wells containing live FhNEJ. Lower asterisks indicate significant differences between the negative control PLG + PAs without FhNEJ and its counterpart wells containing 1 μg of FhNEJ-Teg extract. The differences between the remaining conditions were not statistically significant (*p≤0.05, **p≤0.001, ***p≤0.001; one-way ANOVA followed by Tukey contrasts for pair-wise comparisons).

### 2D analysis of FhNEJ-Teg and identification of PLG-binding proteins by mass spectrometry

In order to determine the identity of the PLG-binding proteins in FhNEJ-Teg, all the proteins of this extract were first separated using 2D electrophoresis spanning a MW range of 15-260 kDa and pI 3-10 (Figure 5A). Secondly, 2D gels were electro-transferred to nitrocellulose membranes in order to detect the presence of PLG-binding proteins by ligand blotting (Figure 5B). By matching the spots in the ligand-blot with those in the homologous silver-stained gel, we identified a total of 33 protein spots that were manually excised from the 2D gel and analyzed by LC-MS/MS. Based on similarity to *F. hepatica* sequences contained in the Uniprot_trematoda database, our proteomic analysis identified an average of 12 proteins per spot and revealed that the tegument of FhNEJ contains 279 potential PLG binding protein isoforms, corresponding to 133 proteins. Figure 6A shows the proteins (including isoforms) that were identified equal to or more than ten times, being the juvenile-specific cathepsin L3 (FhCL3) the most recurrently identified PLG-binding protein in the pool of potential PLG-binding proteins detected by MS in the tegument of FhNEJ. We also identified cathepsins belonging to additional juvenile-specific clades, such as FhCB2 and FhCB3, and PLG receptors identified in other parasite and non-parasite pathogen species, such as enolase, actin, heat-shock protein 70 (hsp70), annexin, glyceraldehyde-3-phosphate dihydrogenase (GAPDH) and fructose-bisphosphate aldolase (FBAL) (Figure 6A and Supplementary Table 1). Gene Ontology annotation of the identified potential PLG-binding proteins revealed that these are mostly involved in biological processes related to proteolysis, regulation of catalytic activity and carbohydrate metabolism (32%, 14% and 12%, respectively) (Figure 6B).

**Figure 5.**
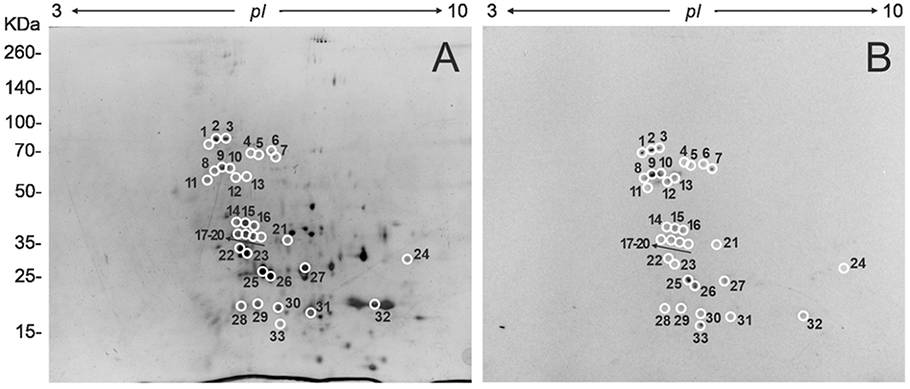
Bidimensional electrophoresis of FhNEJ-Teg and detection of plasminogen binding by immunoblotting. A) 40 μg of FhNEJ-Teg was separated in two dimensions using IPG strips with a pH range of 3-10 and 12% SDS-PAGE gels. B) Immunoblot detection of PLG binding to FhNEJ-Teg proteins. PLG-binding protein spots are circled and numbered.

**Figure 6.**
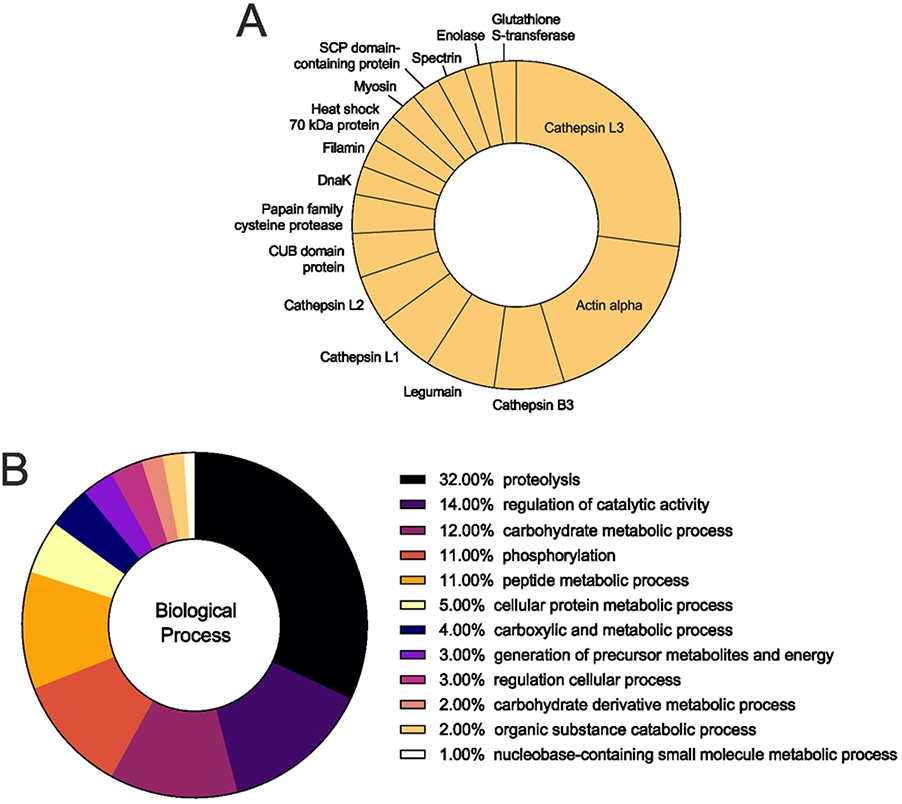
Identification of potential PLG-binding proteins in the tegument of FhNEJ-Teg by mass spectrometry. Protein spots detected in the ligand blot were manually excised and analyzed by liquid chromatography coupled to tandem mass spectrometry (LC-MS/MS). A total of 33 protein spots were identified, corresponding to 133 different proteins. A) Abundance distribution of the 16 most recurrently identified proteins as potential PLG receptors in the tegument of FhNEJ (identified in equal to or more than 10 instances). B) Gene Ontology analysis of the potential PLG-binding proteins identified in the PLG-reactive protein spots.

### Validation of 2D-MS results with recombinant juvenile-specific cathepsins

Our 2D-MS approach showed that juvenile-specific cathepsins B2, B3 and L3 are potentially involved in PLG binding, and we validated these results using recombinant versions of them. The recombinant cathepsins B2, B3 and L3 used for validation had 84.6%, 81.9% and 90.2% identity, respectively, to the sequences identified by MS (data not shown) and were catalytically-inactive (29). Our ELISA-based PLG binding assay confirmed that FhCB2, FhCB3 and FhCL3 are capable of binding PLG in a concentration-dependent manner, being FhCB2 and FhCL3 the proteins with greatest affinity for PLG (Figure 7). PLG binding by these proteases was dramatically reduced in the presence of the lysine analog ε-ACA, which indicates that PLG binding occurs via its kringle domains.

**Figure 7.**
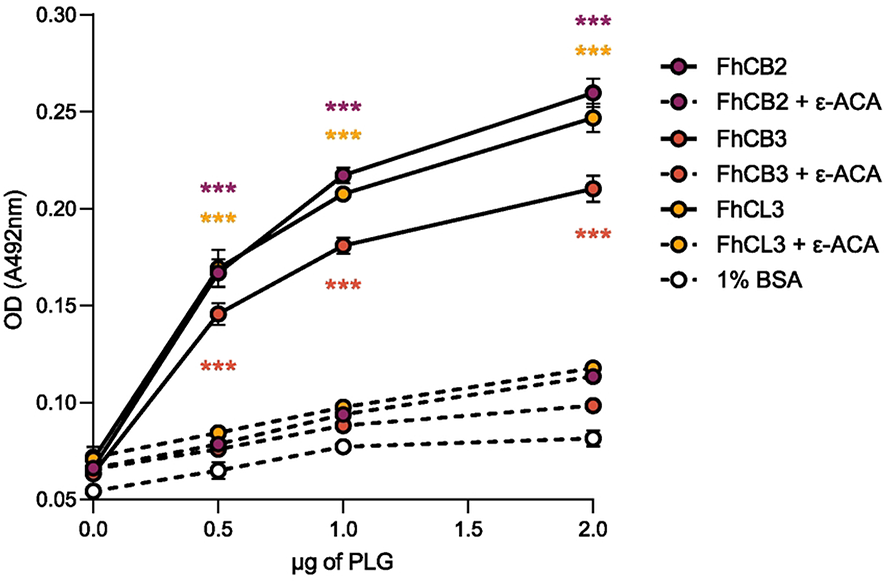
The juvenile-specific cathepsins FhCB2, FhCB3 and FhCL3 bind plasminogen in a concentration- and lysine-dependent manner. The capability of FhNEJ cathepsins B2, B3 and L3 to bind increasing amounts of PLG was assessed via ELISA using recombinant versions of these proteins. In parallel, a competition assay was performed that included 50 mM of ε-ACA during PLG incubation; and incubation with 1% BSA served as a negative control for PLG binding. Dots indicate the mean of three replicates ± SD. Asterisks indicate significant differences between every protein and both its 50mM ε-ACA counterpart condition and the negative control 1% BSA (***p≤0.001; one-way ANOVA followed by Tukey contrasts for pairwise comparisons).

Additionally, we performed plasmin activation enzymatic assays in the presence of recombinant FhCB2, FhCB3 and FhCL3 to address whether binding of PLG by these proteins is functionally relevant in terms of plasmin generation. These assays showed that binding of PLG by FhCB2 and FhCB3 enhances PLG cleavage and conversion to plasmin (Figure 8), although to different extents and requirements in terms of PLG activators. Addition of FhCB2 to the reaction enhances PLG conversion to plasmin preferentially by u-PA over t-PA (Figure 8A), while the opposite holds true when FhCB3 is the protein involved in PLG binding (Figure 8B). Interestingly, and even though Figure 7 shows that FhCL3 is capable of binding PLG, stimulation of bound PLG into plasmin is less potent than that driven by FhCB2 and FhCB3 and only statistically significant at later time points and when t-PA is added to the reaction (Figure 8C).

**Figure 8.**
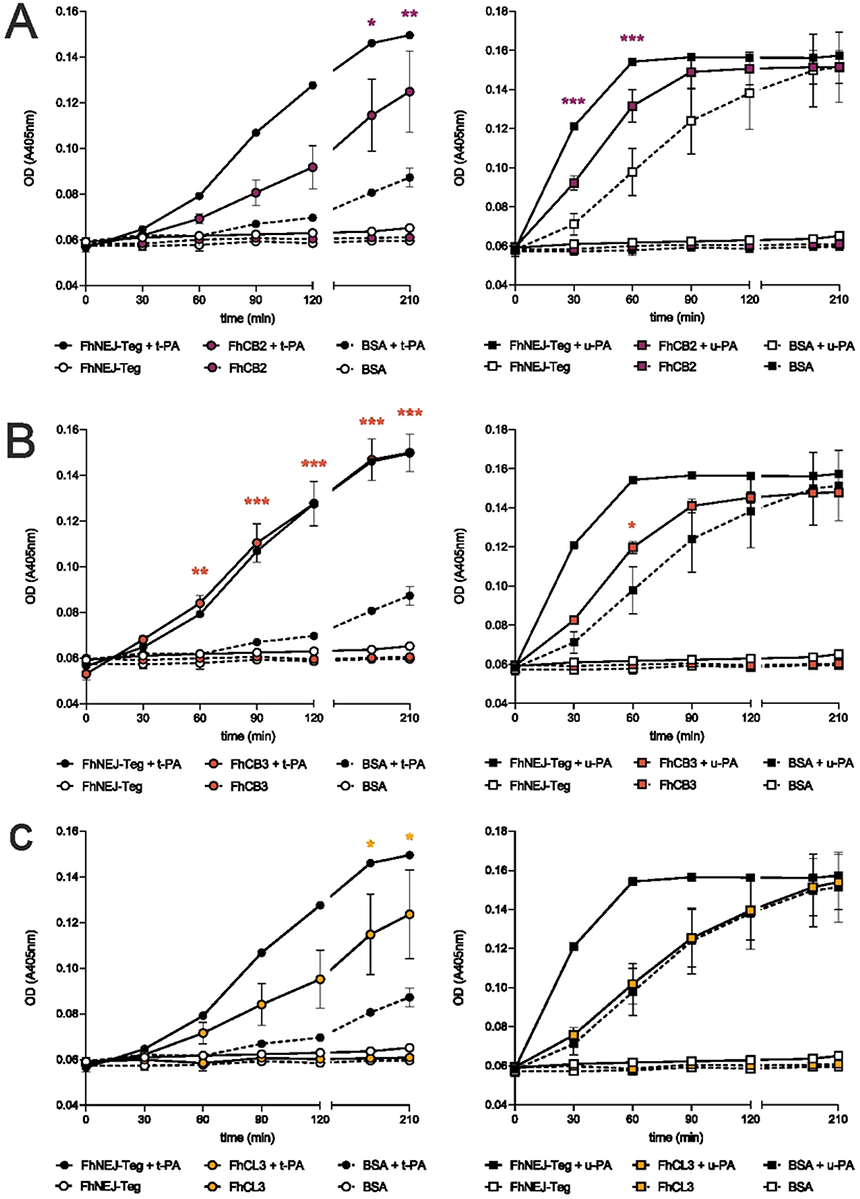
Plasminogen binding by FhNEJ cathepsins enhances plasmin generation. One μg of recombinant FhCB2 (A), FhCB3 (B) or FhCL3 (C) were incubated with human PLG, a chromogenic substrate for plasmin (S-2251) and either t-PA (left panels) or u-PA (right panels) and plasmin generation was assessed by measuring substrate cleavage at OD 405 nm. In some instances, recombinant cathepsins were replaced by 1% BSA as a negative control. Dots indicate the mean of three replicates ± SD, and asterisks indicate significant differences between conditions incubated in the presence of cathepsins + PLG activators and its 1% BSA + PLG activators counterpart (*≤0.05, **p≤0.001, ***p≤0.001; one-way ANOVA followed by Tukey contrasts for pairwise comparisons).

## Discussion

The interaction between parasites and the fibrinolytic system of the host has been demonstrated in very diverse organisms, including bacteria, fungi and parasites, and proposed to be beneficial for parasite survival and migration inside the vertebrate host and also a mechanism of immune evasion (13–15,30). Adult flukes of *F. hepatica* express and secrete proteins that are capable of interacting with PLG and stimulate its conversion to the effector protease, plasmin (16,17), and we hypothesized that this interaction may also occur in the juvenile stages of the parasite and serve as a mechanism for FhNEJ to overcome the intestinal wall and initiate tissue migration throughout the mammalian host.

In order to test whether FhNEJ contain PLG-binding proteins on their tegument surface, we isolated tegument proteins from freshly excysted FhNEJ and analyzed PLG binding by ELISA. This experiment showed that indeed FhNEJ-Teg contains proteins that are capable of binding PLG in a concentration- and lysine-dependent manner. Binding of PLG to its physiologic receptors depends on specific protein domains of the PLG molecule, termed kringle domains, which are capable of binding lysine residues present in partner proteins (18), and we observed that the interaction between FhNEJ-Teg proteins and PLG is also lysine-dependent. The fact that the mechanism of PLG binding to its physiologic receptors is maintained in FhNEJ tegument proteins reinforces the idea that this binding is specific and biologically relevant in the context of tissue invasion and parasite establishment within the mammalian organism.

Complementary to our ELISA assays, we performed immunofluorescent staining of FhNEJ surface-bound PLG to see whether PLG-binding proteins are diffusely expressed throughout the entire tegument surface or specifically located in certain areas. Despite observing some inter-individual variability, we detected a stable pattern of staining at the excretory (protonephridial) pore, which represents a major site of protein secretion together with the tegument and gut (31). In addition to that observed in the excretory pore, some FhNEJ also showed specific PLG labeling at the oral sucker, which together with the ventral sucker is involved in FhNEJ tissue migration via alternate attachment and release (8). This observation further advocates for a role of PLG binding on the surface of FhNEJ in the traversal of the intestinal wall and subsequent FhNEJ tissue migration. We found an additional pattern of PLG staining that was specific not only at the excretory pore and/or the oral sucker but widely spread all over the tegument, which overall suggests that the mechanisms of translocation of PLG-binding proteins to the surface of FhNEJ may be manifold.

Next, we sought to analyze whether PLG binding to the tegument of FhNEJ is functionally relevant by measuring whether it enhances conversion of bound PLG to its catalytically-active form, plasmin. Three hours was considered a biologically relevant time span to measure plasmin generation given that FhNEJ take around two to three hours to cross the intestinal wall after excystment (1), meaning that plasmin generation should occur within this timeframe if it were to be involved in FhNEJ migration through the intestinal epithelium. By incubating FhNEJ-Teg proteins with a chromogenic substrate for plasmin and the PLG activators t-PA or u-PA, we confirmed that binding of PLG by tegument proteins potentiates its conversion to plasmin by both activators. Unlike described in *Schistosoma bovis*, a closely-related trematode species (32), we were unable to detect any plasmin activity in the absence of PLG activators, indicating that the tegument of FhNEJ does not contain proteases that are capable of directly activating PLG. We envision that the mechanism behind enhanced PLG activation in the presence of PLG-binding tegument proteins is most likely similar to that described for canonical PLG receptors, i.e., through the induction of a conformational change in bound PLG molecules that exposes cleavage sites for t-PA and/or u-PA, thus facilitating the generation of plasmin (19,33). Remarkably, when FhNEJ-Teg extract was replaced by an equivalent amount of live FhNEJ, we confirmed that PLG binding by FhNEJ enhances plasmin generation in the presence of PLG activators. Given that these FhNEJ remained alive throughout the entire experiment, we cannot rule out the possibility that plasmin generation in non-tegument surfaces, including the gut, or by FhNEJ excreted/secreted products, as observed in adult *F. hepatica* (17), could also account for the enhancement of plasmin generation observed in wells containing live FhNEJ. In relation to this, a recent publication using laser microdissection to generate exclusive tissue fractions of the tegument and the gut of adult *F. hepatica* showed that detergent-based extraction of *F. hepatica* tegument antigens may also enrich this antigenic fraction with proteins derived from the parasite’s gastrodermal cells (34). Therefore, it still remains a possibility that tegument PLG-binding proteins may have arisen from gut secretions and adhered to the parasite’s surface during incubation prior to fixation (12). Notwithstanding the origin of the tegument PLG-binding proteins, and based on the capacity of plasmin to degrade components of the ECM (35), these results support our original idea that PLG fixation on the tegument surface could represent a mechanism for FhNEJ to harness the functions of the fibrinolytic system that provides an additional source of proteolytic activity to facilitate traversal of the intestinal wall and the successful establishment of the parasite within the vertebrate host.

The PLG activation/plasmin generation activities of t-PA and u-PA are related to variable physiological processes due to differential tissue expression of these proteases. Whereas t-PA expression is almost restricted to the vascular endothelium (18,19), u-PA is synthesized by a wider range of cell types, such as endothelial cells, macrophages, neutrophils, renal epithelial cells, some tumor cells (18,36) and epithelial cells of the small intestine (37). As a result, t-PA activity is mostly restricted to intra-vascular fibrinolysis (19); whereas u-PA is considered a master driver of cell migration (38) in different physiological settings, namely vascular remodeling/angiogenesis, tumor invasion and the inflammatory response (19,39). The observation that plasmin generation seems to be preferentially enhanced when u-PA rather than t-PA is added to the reaction is most likely explained by the fact that u-PA is capable of partially activating soluble (unbound) PLG whereas t-PA is not (33). Despite this technical constraint, we presume that u-PA may be the activator involved in plasmin generation from FhNEJ surface-bound PLG in a real physiological setting given that this is the PLG activator that is most highly expressed in the small intestine (37).

Aiming at identifying the proteins that are responsible for PLG binding in the tegument of FhNEJ, we performed a proteomic analysis using 2D electrophoresis and ligand blotting coupled to mass spectrometry that identified 133 proteins potentially capable of binding PLG (summarized in Supplementary Table 1). We partly validated these results by performing an ELISA-based PLG binding assay with recombinant versions of cathepsins B2, B3 and L3, whose presence at the FhNEJ tegument surface has been described (12), but we envision that the majority of the proteins identified by 2D-MS as potential PLG receptors might indeed be used for PLG binding purposes in a real physiological setting. It is not surprising that a large number of proteins are used by FhNEJ for PLG recruitment provided that in pathogenic organisms, including parasites, PLG binding and plasmin activation is also highly redundant since multiple surface receptors normally coexist for this purpose (13–15). From an evolutionary perspective, redundancy is an efficient strategy to ensure the maintenance of essential biological functions since it provides for back-up proteins to take over in case of loss of function of other effector proteins due to genetic mutations (40). Given that utilizing proteins for one single and same (redundant) function would result in genetic drift and subsequent loss of redundancy, a successful strategy to preserve redundancy is to employ proteins that have both overlapping (redundant) and unique (non-redundant) and independent functions so that selective pressure in favor of the proteins’ independent roles guarantees the maintenance of their expression over time (41). This is precisely what we observe in this study: the use of proteins with highly divergent biological functions for PLG binding/plasmin activation purposes (see Figure 6A for a GO annotation of the potential PLG-binding proteins identified by 2D-MS), which reinforces the notion that the interaction between FhNEJ and the fibrinolytic system of the host ensures parasite fitness and thus represents an essential mechanism for parasite survival inside the mammalian organism. Unfortunately, the fact that FhNEJ interact with the host fibrinolytic system via such redundant mechanisms poses great difficulties to confront this interaction through classic control and therapeutic strategies (13).

Among others, we identify some of the best characterized PLG receptors in different bacterial, fungal and parasite species as potential PLG-binding proteins in the surface of FhNEJ, such as GAPDH, FBAL, enolase and annexin (14,15,30). These proteins are also present in the surface of certain tumor cells and their over-expression is linked to metastasis (42), further highlighting the role that the PLG/plasmin system plays in diverse biological processes in which cell migration is required. Interestingly, some of these canonical PLG receptors, including enolase, GAPDH and FBAL, are glycolytic enzymes devoid of transmembrane domains and hence their classical biological role is cytosolic. In fact, the third most represented biological function identified in the pool of PLG-binding proteins in FhNEJ-Teg is carbohydrate metabolism, which is not surprising considering that the use of glycolytic enzymes for PLG binding purposes has been well documented in many pathogenic and non-pathogenic organisms, including multicellular parasites (14). The mechanism by which FhNEJ GAPDH, FBAL and enolase are transported to the tegument surface, the outermost layer of FhNEJ, still remains an open question. In other organisms, several mechanisms for cell surface expression of these proteins have been proposed, including relocation to the plasma membrane as part of a protein complex, binding to lipid molecules followed by translocation to the outer membrane, and non-covalent association to the cell membrane after active secretion to the extracellular space (14,42). Alternatively, these proteins could also be released to the extracellular space as part of the cargo of extracellular vesicles (EVs), which have recently been identified in FhNEJ (43) and proposed to represent the secretion route of *F. hepatica* molecules that lack consensus N-terminal signal sequences for their secretion via the ER/Golgi pathway (12). In line with this, a proteomic analysis of FhNEJ EVs revealed that these secretory structures harbor glycolytic enzymes identified in the present study as potential PLG-binding proteins, namely GAPDH (Accession A0A2H1BWY6), heat-shock protein 70 (Accession B1NI98), heat-shock protein 90 (Accession A0A2H1BZF7) and CUB-domain protein (Accession A0A2H1C1S8) (44). Additionally, adult *F. hepatica* smallest EVs (the so-called 120K EVs) harbor GAPDH (45), and another report demonstrates that these 120K EVs are most likely released via the protonephridial (excretory) system (31). Although we do not know whether FhNEJ smallest or 120K-like EVs are the ones carrying GAPDH (and/or other PLG-binding proteins identified in this study) since the proteome of FhNEJ EVs has so far been analyzed in total FhNEJ EVs and not separate populations (44), these observations coincide with the staining pattern of PLG binding observed in our immunofluorescence images, which is most prominent in the protonephridial (excretory) pore, further supporting the idea that some of the PLG-binding proteins that we detected on the surface of FhNEJ may be released as part of the cargo of FhNEJ EVs.

In addition to glycolytic enzymes, proteolysis is the most represented biological function of the potential PLG-binding proteins identified in our 2D-MS analysis. Remarkably, the most recurrently identified potential PLG-binding proteins belong to the superfamily of cathepsin proteases. More specifically, we identify FhCL3 and FhCB3 as potential PLG-binding proteins, and FhCL1, FhCL2 and FhCB3 to a lesser extent. This is consistent with the developmental stage-specific expression of these proteases, being FhCB1-3 and FhCL3 mostly expressed in the juvenile stages of the parasite that are found in the duodenum (46–48). However, one of the limitations of our 2D-MS approach is that it is performed under denaturing conditions, which may expose internal PLG-binding epitopes in some proteins that are not employed for PLG-binding in a real physiological context. Therefore, we set out to validate our 2D-MS results by confirming PLG binding in an ELISA experiment, which is performed under native conditions, using recombinant and catalytically-inactive versions of the juvenile cathepsins B2, B3 and L3 (29). This experiment confirmed the capacity of these proteins to bind PLG in a concentration- and lysine-dependent manner, and our plasmin activation enzymatic assays confirmed that FhCB2 and FhCB3 also serve as enhancers of plasmin generation from bound PLG, although with different kinetics and involving different PLG activators. Whereas FhCB2 induces plasmin generation by u-PA and not t-PA, the opposite holds true for FhCB3, and even though we confirmed that FhCL3 is capable of binding PLG, this protein is apparently less effective in stimulating its conversion to plasmin.

Previous experiments performed in our lab following a similar 2D-MS strategy showed that cathepsins present in excretory/secretory products of *F. hepatica* adult flukes are capable of binding PLG (17), and we now show that the FhNEJ-specific cathepsins B2, B3 and L3 behave likewise. To the best of our knowledge, *F. hepatica* is the only parasite whose cathepsins have been identified as PLG-binding proteins (13), and given that we validated our results using catalytically-inactive recombinant versions of them, we can also conclude that this is the first time that a proteolysis-independent function is described for these proteases in relation to FhNEJ migration. Therefore, and based on the present results, we propose that *F. hepatica* cathepsins shall be regarded as moonlighting proteins, which are defined by their capability to play two or more unrelated roles depending on their cellular or developmental context (49). Proteins that are constitutively expressed and have high levels of expression are more likely to develop moonlighting functions because their constitutive nature makes it more likely for them to be involved in advantageous interactions with protein and non-protein partners. Similarly, proteins that arise from gene duplication events are likely to moonlight given that gene duplication creates a permissive space in which new functions can be developed without compromising the canonical role of the protein (49). *F. hepatica* cathepsin peptidases are amongst the most abundantly expressed proteins in FhNEJ (50) and they are also products of gene duplication events (51), which makes them good candidates to acquire moonlighting roles including the PLG-binding function that we describe in this study.

From a clinical perspective, the interaction between FhNEJ and the fibrinolytic system described in this study may contribute to and/or exacerbate focal hemorrhages that are caused by migratory juveniles during the acute phase of infection (1,52). Furthermore, our results could also be relevant in the context of neurological manifestations associated to this disease, provided that minor neurological manifestations including cephalalgias, character disorders, vertigo, nightmares, delusional disorders, and/or insomnia are not rare (53) and they have been proposed to be mediated by increased blood-brain barrier permeability caused by an aberrant fibrinolytic balance in infected patients (17). These neurological symptoms preferentially occur during the acute phase of infection (53) and we now show that FhNEJ are capable of interacting with the host’s fibrinolytic system, thus giving a plausible mechanistic explanation to the appearance of the aforementioned neurological disorders during acute fasciolosis.

In conclusion, we have demonstrated for the first time that FhNEJ interact with the fibrinolytic system of the mammalian host as a potential early stage invasion mechanism by binding PLG and enhancing its conversion into the active enzyme, plasmin. We further determined that PLG binding by FhNEJ is highly redundant and mainly driven by juvenile cathepsin peptidases, which advocates for a previously uncharacterized proteolytic-independent role of these enzymes in relation to FhNEJ trans-intestinal migration. Altogether, this work contributes to a deeper understanding of host-parasite relationships in the earliest phases of fasciolosis, and sheds light into a potential FhNEJ migration mechanism that may be worth exploiting from a pharmacological and immunological perspective for the development of more successful treatment and control strategies against this widespread parasitic disease.

---

**Supplementary Table 1. Potential PLG-binding proteins in the tegument of FhNEJ identified 2D-LC-MS/MS.**

## Acknowledgements

We thank Prof. John P. Dalton (University of Galway) for the recombinant constructs used in our validation experiments, K. Cwiklinski (University of Liverpool) and N.E.D. Calvani (University of Galway) for their fruitful tips on immunofluorescent staining of FhNEJ and C. Castro (Institute of Functional Biology and Genomics, CSIC/USAL) for her support during the acquisition of confocal images.

